# In vivo characterization of Achilles subtendon function and morphology within the tendon cross section and along the free tendon

**DOI:** 10.1101/2025.05.29.656873

**Authors:** Kathryn S. Strand, Todd J. Hullfish, Josh R. Baxter

**Affiliations:** Department of Orthopaedic Surgery, University of Pennsylvania, Philadelphia, PA, USA

**Keywords:** Achilles Subtendons, ultrasound, muscular stimulation

## Abstract

The Achilles tendon is composed of three distinct fascicle bundles, or “subtendons,” each originating from the head of one of the three triceps surae muscles. In a healthy tendon, these subtendons slide relative to each other during muscle contractions. This subtendon sliding is reduced in older adults and younger adults who suffer an Achilles tendon injury. However, subtendon sliding is challenging to quantify in low-load scenarios that are critical for monitoring subtendon biomechanics in patients with mechanically compromised tendons, like following an Achilles tendon rupture and repair surgery. The purpose of this study was to develop a reliable method to characterize subtendon behavior *in vivo* using combined transverse plane ultrasound imaging and neuromuscular electrical stimulation of individual gastrocnemii. We used a Kanade-Lucas-Tomasi point tracking algorithm to quantify tendon displacement during isolated muscle stimulations. Next, we applied k-means clustering to characterize heterogeneous subtendon behavior within the tendon cross section. The tendon cross section displayed differential displacement patterns depending on the stimulated muscle (p<0.0001), and these displacements differed along the free tendon during lateral gastrocnemius stimulations (p=0.004). These results reflect possible differences in load-sharing between adjacent subtendons and differing muscle-tendon dynamics among the triceps surae muscles. Finally, this method confirmed bilaterally symmetric subtendon behavior and demonstrated high inter-session reliability (ICC>0.83). Overall, this study furthers our understanding of differential muscle-tendon dynamics of individual Achilles subtendons both within the tendon cross section and along the free tendon. Future work will apply this method to injured populations to develop biomarkers of altered subtendon function.

**NEW & NOTEWORTHY:** Achilles subtendon function and morphology are challenging to characterize *in vivo*. This study employed transverse plane ultrasound imaging and neuromuscular electrical stimulation to characterize behavior of individual subtendons both within the tendon cross section and along the free tendon. It is the first study to demonstrate functional behavior within the Achilles tendon using these combined tools in identifying subtendon dynamics. Additional findings of bilateral symmetry in healthy individuals present this tool’s potential to quantify altered subtendon function post-injury.

## Introduction

The Achilles tendon is a complex structure composed of three distinct fascicle bundles, or “subtendons,” that originate from the medial gastrocnemius (GM), lateral gastrocnemius (GL), and soleus muscles (3,4). These subtendons twist such that the medial portion of the tendon inserts posterior, and the lateral portion inserts anterior into the calcaneus (5,6). Additionally, these subtendons slide relative to each other in young adults with healthy tendons which is typically quantified by non-uniform displacements of superficial and deep tendon regions detected via ultrasound imaging during plantar flexion (1,2). In contrast, subtendon sliding is reduced in an injured tendon (7–10). For example, Beyer et al. found that the deep Achilles tendon layer displaced 50% more than the superficial Achilles tendon layer, but subtendon sliding was not detected in Achilles tendons that were healed following a rupture repair surgery (10). Similarly, Lecompte et al. found that superficial and deeper layers of healthy Achilles tendons experienced an average of 78% more relative sliding compared to tendinopathic tendons (8). Classic studies on plantar flexor muscle morphology in trained sprinters highlight the importance of subtendon sliding to efficiently transfer loads from muscles with different force-length properties (11). Conversely, decreased plantar flexor function in adults recovering from Achilles tendon injuries may be constrained by an Achilles tendon that has more uniform displacements. These functional changes highlight the need to develop reliable tools to quantify subtendon sliding to test the effects of training, injury, and aging on Achilles tendon biomechanics and plantar flexor function.

*In vivo* characterization remains a clinical challenge due to the limitations of common imaging modalities despite the growing interest in Achilles subtendon structure and function. A study by Cone et al. utilized high-field (7-Tesla) magnetic resonance imaging (MRI) to identify subtendon boundaries (12). While effective in characterizing three-dimensional tendon structure, this imaging modality is impractical in most research and clinical settings. Ultrasound imaging has been used extensively by the biomechanics community to characterize the dynamic behavior of the Achilles tendon with recent efforts aimed at elucidating load-sharing behavior between adjacent subtendons. For example, both Klaiber et al. and Finni et al. used simultaneous transverse plane ultrasound imaging and neuromuscular electrical stimulation (NMES) to estimate subtendon morphology based on localized displacements within the tendon cross section in response to contractions of individual triceps surae muscles (13,14). NMES is advantageous over voluntary contractions because it ensures that isolated muscle recruitment corresponds to observed tendon behavior. This isolated recruitment also allows for investigation into the passive load sharing between adjacent subtendons due to the interfascicular matrix (15,16).

Prior studies combining transverse plane ultrasound imaging and NMES solely evaluate Achilles tendon morphology but do not quantify functional differences between subtendons. Many studies characterize subtendon function using ultrasound imaging in the coronal or sagittal planes to quantify the displacements of the superficial and deep layers of the tendon (2,17–20). Other studies quantifying tendon motion in the transverse plane have done so to classify Achilles tendon twist (13,14) rather than describe subtendon function. Although the tendon is loaded axially, load sharing between subtendons may also manifest transversely. For example, intratendinous pressure that results from the torsion of the 3 subtendons may be a driver of tendinopathy (21). The scar tissue that forms in a ruptured tendon may increase interfascicular adhesions and reduce sliding between subtendons. Moreover, prior studies limit tendon evaluation to a single imaging plane, so there is still a need to evaluate dynamic *in vivo* Achilles tendon behavior in multiple dimensions. Doing so is particularly important for studying Achilles tendon injuries like tendinopathy and rupture that are characterized by heterogenous tissue composition.

The purpose of this study was to develop a reliable *in vivo* method to characterize Achilles subtendon function and morphology along the length of the free tendon in healthy subjects. We used simultaneous transverse plane ultrasound imaging and NMES to individual triceps surae muscles and quantified localized tendon movement using an automated point tracking algorithm. We hypothesized that 1) the tendon would demonstrate heterogenous behavior in response to isolated muscle stimulations along its length and cross section, 2) subtendon behavior would display bilateral symmetry, and 3) this method would reliably (ICC>0.67) (22) identify localized regions of tendon displacement across separate testing sessions.

## Methods

Fifteen healthy adult subjects (7M, 8F, age 25±2 years, 10 Caucasian/White, 4 Asian, 1 African American/Black) with no history of Achilles tendon injuries or neuromuscular deficits provided informed, written consent to participate in this study approved by our University’s Institutional Review Board. A priori power analysis indicated that this sample size was sufficient to achieve a power of 0.96 (effect size: 1.13, alpha=0.05) calculated from a pilot study of n=8 subjects based on the hypothesized bilateral symmetry. Subjects lay prone on a bed with their legs fully extended and the ankle in a neutral, relaxed position. We fixed a 21 MHz linear ultrasound transducer (L6-24, LOGIQ E10, GE HealthCare, Chicago, IL, United States) perpendicular to the free tendon with a custom 3D printed fixture to acquire ultrasound images in the transverse plane. Imaging parameters were as follows: gain: 45, dynamic range: 63, depth: 2-3 cm. Similar to previous studies (13,14,18), we delivered electrical stimulations via hydrogel electrode pairs (25×38 mm) placed over the muscle bellies of the GM and GL muscles (inter-electrode distance: 2 cm) (**Figure 1a**). We placed additional electromyography (EMG) recoding electrode pairs (12×10 mm) over each gastrocnemius and the medial portion of the soleus muscle, distal to the GM, as well as reference and grounding electrodes over the medial and lateral malleoli. We delivered monophasic pulse trains (100 Hz, 400 µs pulse width, 1 s duration) using a constant current stimulator (DS8R, Digitimer Ltd, Hertfordshire, United Kingdom) beginning at 4 mA and increasing by 2 mA until there was visible plantar flexion of the foot. Once the stimulation amplitude induced plantar flexion, we decreased the stimulation amplitude by 1-2 mA such that there was no visible plantar flexion, but localized tendon displacement remained visible in the ultrasound video. This method ensured isolated recruitment of the stimulated muscle that we confirmed using EMG measurements. We then placed the ultrasound probe 1 cm proximal to the medial malleolus and delivered stimulations to the GM and GL muscles at their respective stimulation amplitudes. Next, we moved the probe 1 cm proximally along the free tendon toward the muscle-tendon junction and repeated the muscle stimulations. We repeated these steps on both legs at 5 locations along the free tendon (**Figure 1b**). As the free tendon is approximately 4-7 cm long (23), these increments ensured imaging of the entire tendon.

**Figure 1.**
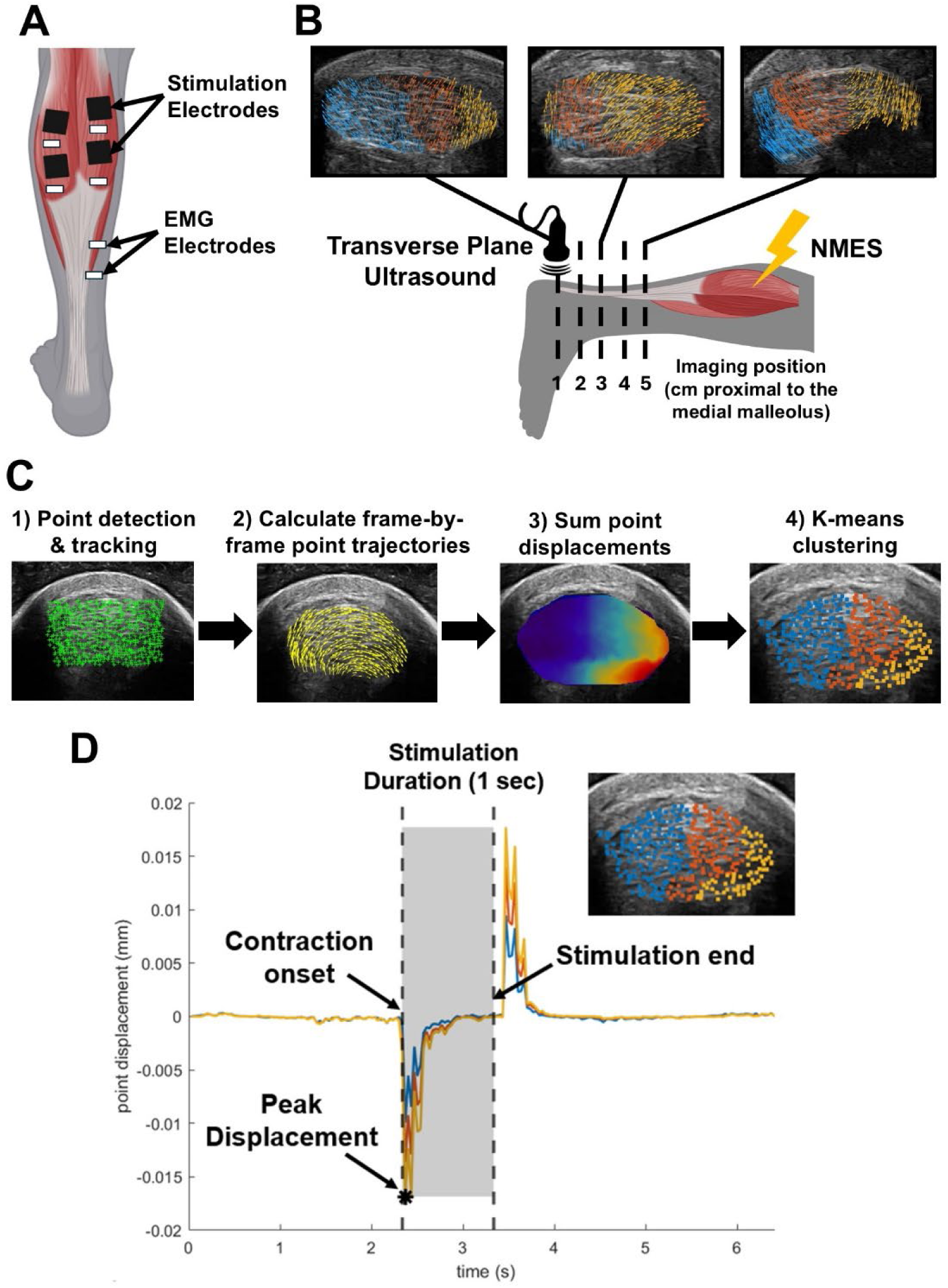
Experimental setup. A) Placement of NMES stimulation electrodes and EMG recording electrodes. B) Ultrasound imaging setup with ankle in a neutral position and transverse ultrasound imaging at five locations along the free tendon. Vector fields overlaid on ultrasound images of the Achilles tendon cross section demonstrate heterogeneous responses of the tendon to NMES along the free tendon. C) Point tracking and k-means clustering workflow. D) Example mean trajectories of points within the identified clusters. Displacement values rapidly increase upon stimulation onset and rapidly decrease after the stimulation ends and the tissue relaxes.

We chose to only stimulate the gastrocnemius muscles, as pilot testing revealed that GL and GM stimulations resulted in negligible co-activation of the soleus. Exclusion of the soleus also allowed us to image past the soleus muscle-tendon junction to observe the gastrocnemii in more isolation. However, future work will expand this method to include soleus stimulations and images at more distal regions of the free AT. The mean stimulation amplitude for the GL and GM muscles was 12.2±5.3 mA and 10.3±4.3 mA, respectively. We confirmed that this NMES paradigm isolated the target muscle based on EMG and ultrasound analyses. We found that surface EMG measurements were small in the adjacent muscles (**Table 1**) and these measurements were likely caused by stimulation artifact. We also performed ultrasound imaging of both the stimulated and non-stimulated muscles and quantified fascicle shortening using a previously validated automated tracking algorithm in a subset of participants (n=3) (24) (**Figure S1**). We observed approximately 5 mm of fascicle shortening in the stimulated muscle compared to 2 mm of shortening in the non-stimulated muscle. Prior studies observed a minimum of 4-6 mm of gastrocnemius fascicle shortening during submaximal contractions (25). These findings demonstrated minimal fascicle shortening in the non-stimulated muscle and confirm that our stimulation paradigm produced isolated gastrocnemius contractions.

**Table 1.** Peak RMS values of non-stimulated triceps surae muscles during NMES of the GL and GM.

|  | GL Peak Root-Mean Square (mV) | GM Peak Root-mean Square (mV) | Soleus Peak Root-Mean Square (mV) |
| --- | --- | --- | --- |
| GL Stimulations | --- | 2.29±1.40 | 0.49±0.68 |
| GM Stimulations | 2.90±1.55 | --- | 1.06±0.85 |

We performed a similar protocol on a subset of participants (n=8, 4M, 4F) to establish the repeatability of this method at two clinically relevant ankle positions: the ankle fixed in neutral under an approximate Achilles tendon load of 370 N (26) and the ankle supported in 20 degrees of plantar flexion. We tested these two joint postures to establish reliability in a nominally loaded and unloaded setting. This is important for determining the best testing protocol for future studies that evaluate subtendon mechanics in patients recovering from Achilles surgical treatments that are not cleared for full weight bearing. Subjects lay prone on a dynamometer with their legs fully extended. We fixed the ultrasound transducer perpendicular to the free tendon 3 cm proximal to the medial malleolus. The stimulation amplitude was chosen using the methods described previously with the ankle first fixed in a neutral position, then with the foot fixed in 20° plantar flexion at the same stimulation amplitude. We chose 20° planter flexion to simulate early weight bearing conditions following rupture or surgical intervention when the ankle would be immobilized in plantar flexion to minimize tendon load (27,28). We confirmed passive ankle moments using the dynamometer and found that fixing the ankle in neutral position loaded the ankle with 0.939±1.726 Nm, while supporting the ankle in 20° plantar flexion reduced this load to 0.495±0.409 Nm during stimulations. We plan to use this tool to evaluate tendon healing outcomes in patients following surgical repairs and debridement of Achilles tendon injuries, so minimizing the amount of applied tendon load is an important criterion. We estimate that supporting the ankle in neutral passively loads the Achilles tendon (assuming a 5 cm moment arm (29)) with approximately 14 N of load, which is 95% less than the mechanical strength of suture used to surgically repair the tendon (30). A single investigator performed this protocol on both legs and repeated on a separate day (<5 days later).

We processed experimental data using custom scripts (MATLAB R2024b, MathWorks Inc. Natick, Massachusetts, United States) to quantify tendon movement and automatically detect subtendon regions during each stimulation. First, we selected the series of frames corresponding to the 1 second stimulation. We manually selected the tendon cross section as the tracking region of interest from the first frame in the series. Next, we used a Kanade-Lucas-Tomasi point tracking algorithm (31,32) to identify the displacement of 900 corner point eigenfeatures within this region of interest (**Figure 1c**) (Pyramid Level: 1, Block Size: 191). During preliminary testing, we determined that 900 eigenfeatures was appropriate to track all regions of the tendon cross section, accounting for hypoechoic regions and points lost during tracking due to out-of-plane motion. We chose point tracking over other common methods of ultrasound motion analysis such as speckle tracking which is more sensitive to out-of-plane motion (33). We summed point displacement magnitudes across all frames of the stimulation and performed k-means clustering (k=3 clusters, based on 3 known subtendons) of the points based on this cumulative displacement during the stimulation. Next, we refined these clusters using a density-based clustering algorithm (Minimum Points: 20, Epsilon: 35) to isolate cohesive regions of points and selected the cluster containing the greatest mean cumulative displacement for further analysis. Due to the high probe frequency (21 MHz), each pixel within the images measured approximately 0.04 mm, allowing us to detect sub-millimeter displacements within the tissue. All reported results analyzing subtendon behavior refer to the cluster experiencing the greatest cumulative point displacement. To identity whether these clusters represented subtendon morphology, we calculated the centroids and areas formed by these clusters, as described by previous work (13,14). We represented point movement as heterogeneous vector fields over the tendon cross section, with each vector representing the magnitude and direction of a point at any given frame. To evaluate the tendon’s functional response to each stimulation, we calculated the mean vector angle of the points within the cluster of interest at the frame during which the tendon experienced peak displacement (**Figure 1d**). To control for differences in ultrasound probe orientation in the transverse plane, we fit an ellipse to the tendon region of interest and calculated all point cluster positions and displacement vectors relative to the major and minor axis, representing the medial-lateral and anterior-posterior directions, respectively. Each frame was approximately 37 ms. As a result, our sampling rate was too low to capture the electromechanical delay between stimulation onset and muscle contraction (34), so tissue response to stimulations represents the time between contraction onset and the end of the pulse train.

### Statistical Analysis

To test hypothesis 1, (heterogeneous behavior within the cross section and along the free tendon), we used a two-way repeated measures ANOVA to analyze the effect of stimulated muscle and imaging position along the free tendon on the relative location and area of the selected point cluster as well as the displacement direction within the tendon cross section. Significant main effects and interactions were further analyzed using post-hoc pairwise comparisons with a Bonferroni correction. Differing cluster positions and directional displacement along the free tendon would reflect the AT’s twisted morphology. To analyze potential differences in subtendon sliding along the free tendon, we compared mean cumulative point displacement during both GL and GM stimulations across all five imaging positions along the free tendon using a repeated measures analysis of variance (ANOVA) with a Bonferroni post-hoc correction. Greater cumulative displacement would indicate a higher degree of independent subtendon sliding, while lower values would suggest more interfascicular connections between adjacent subtendons. To test hypothesis 2 (bilateral symmetry), we used paired t-tests to compare cluster positions and areas between the right and left legs. Similar cluster regions on both legs would indicate comparable subtendon responses to muscle stimulation. Finally, to answer hypothesis 3 and establish inter-session measurement reliability of this protocol, we calculated the intraclass correlation (ICC) coefficients with a two-way random-effects model (35) for outcome measures from both joint positions. Significance levels were set to p<0.05 for all tests.

## Results

### Subtendon Kinematics

The Achilles tendon experienced differential displacement patterns depending on the stimulated muscle and imaging position along the free tendon. There was a main effect of stimulated muscle on peak displacement direction (p<0.0001) and an interaction effect between stimulated muscle and imaging position on this outcome (p=0.0035) (**Figure 2**). Post-hoc testing revealed significant differences between displacement direction due to GM and GL stimulations at imaging positions 2 (p=0.046), 3 (p<0.0001), 4 (p=0.0009), and 5 (p<0.0001), as well as differences between positions 1 and 5 during GL stimulations (p=0.0184). During GL stimulations, cumulative displacement at position 1 was significantly less than at positions 3 (p=0.0011), 4 (p=0.0045), and 5 (p=0.0004) (**Figure 3a**). There were no differences in cumulative displacements along the tendon during GM stimulations (**Figure 3b**).

**Figure 2.**
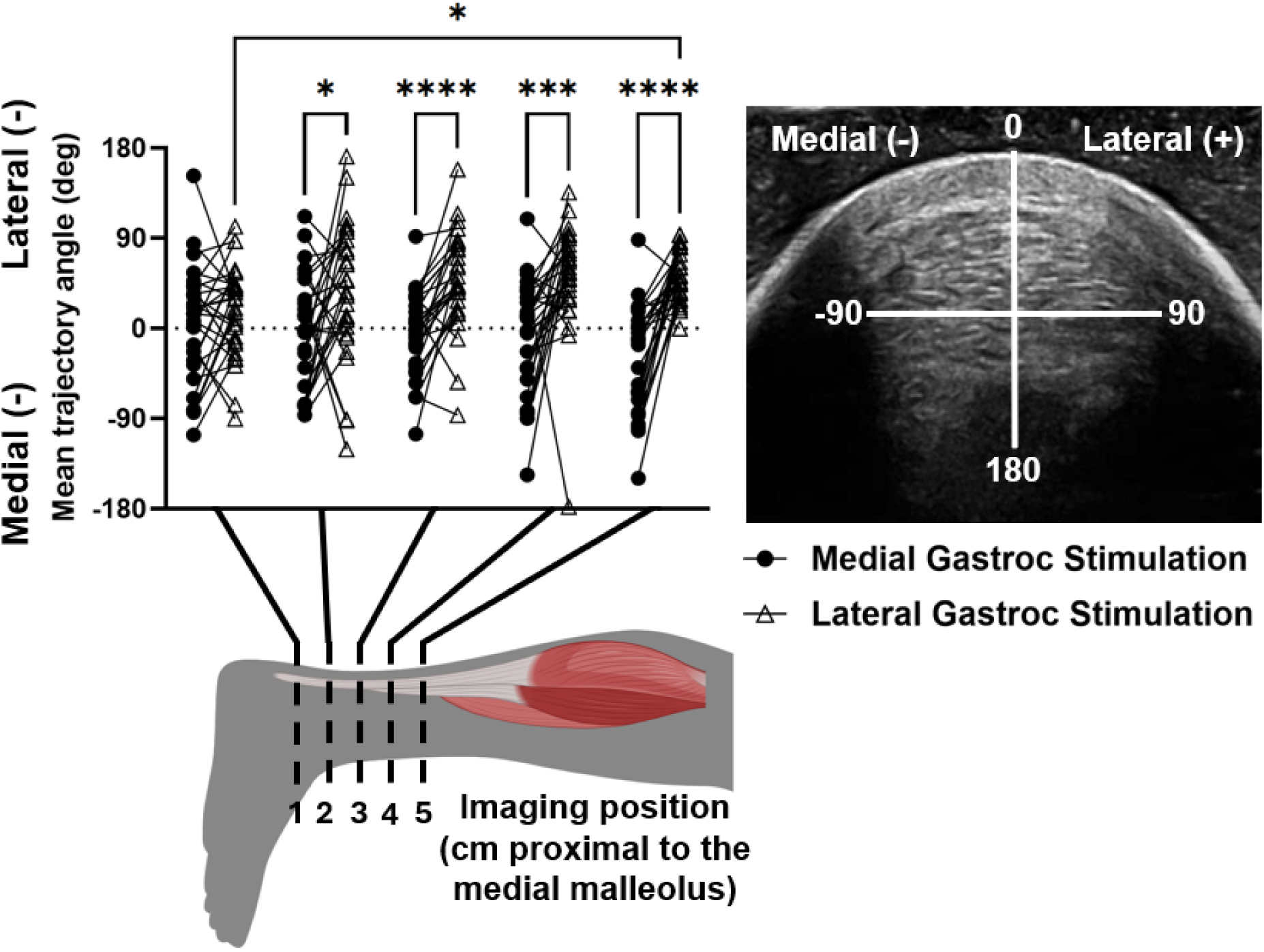
Mean direction of peak displacement of clusters during GM and GL stimulations. *p<0.05, ***p<0.001, ****p<0.0001.

**Figure 3.**
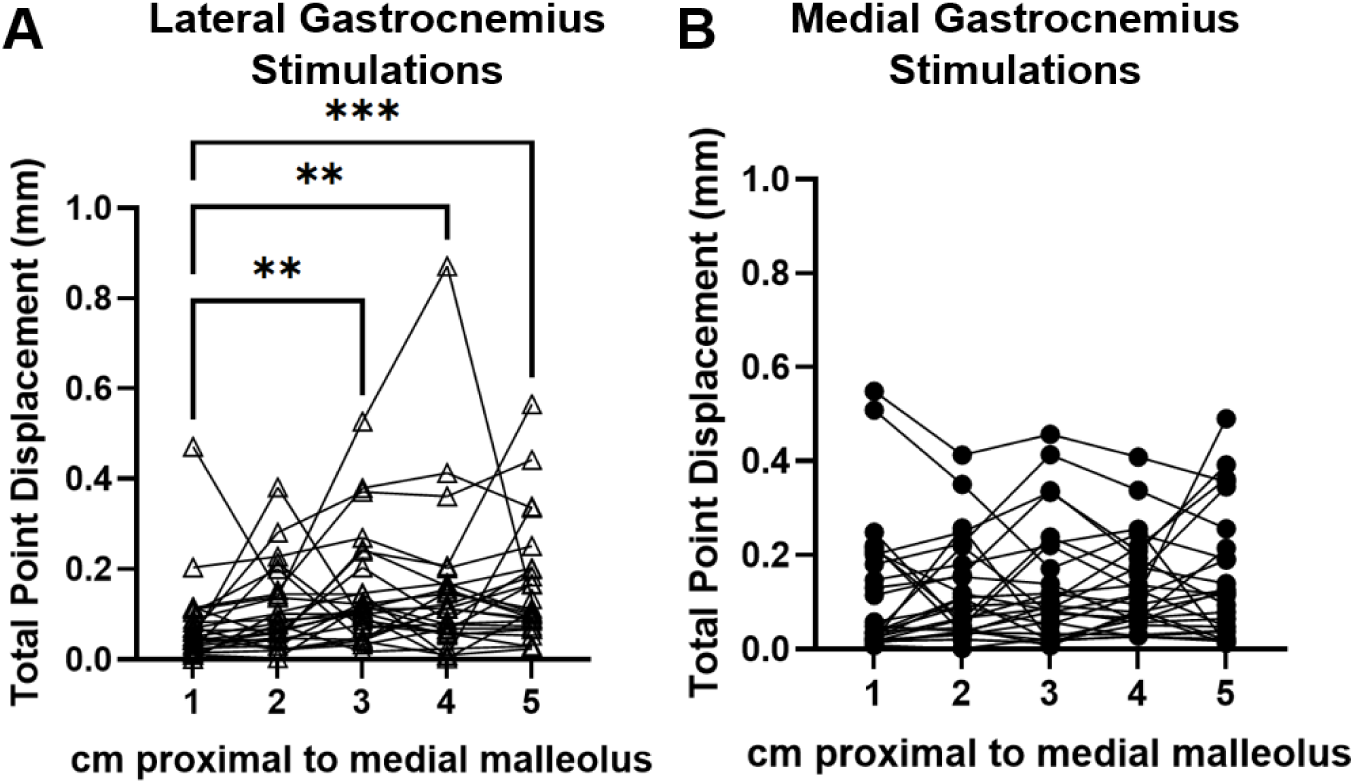
Mean cumulative displacement of points during. A) GL stimulations and B) GM stimulations. **p<0.01, ***p<0.001.

### Subtendon Morphology

Stimulations of GM and GL muscles displaced differing regions of the tendon cross section along the free tendon. There was a main effect of stimulated muscle on cluster medial-lateral position (p<0.0001) and an interaction effect between stimulated muscle and imaging position along the free tendon (p<0.0001). Post-hoc comparisons revealed differences between cluster medial-lateral positions at imaging positions 3 (p=0.0007), 4 (p<0.0001), and 5 (p<0.0001) (**Figure 4a**). During GM stimulations, cluster positions differed between position 1 and positions 3 (p= 0.0062), 4 (p=0.0021), and 5 (p=0.0009), as well as between position 2 and positions 3 (p= 0.036), 4 (p=0.025), and 5 (p=0.0037) (**Figure 4c**). During GL stimulations, cluster medial-lateral positions differed between positions 1 and 3 (p=0.0402), 1 and 5 (p=0.0182), and 2 and 5 (p=0.0461) (**Figure 4d**). There was also a main effect of stimulated muscle on cluster anterior-posterior position (p<0.0001), with differences at imaging positions 1 (p=0.0271), 3 (p=0.0197), and 4 (p=0.0273) (**Figure 4b**). Lastly, there was a main effect of imaging position along the free tendon on cluster area fraction (p=0.0381), with differences between positions 1 and 5 (p=0.0201), 2 and 5 (p=0.0221), and 4 and 5 (p=0.0294) during GM stimulations (**Figure 5**). The medial gastrocnemius cluster was 33.3% and the lateral gastrocnemius cluster was 31.6% of the Achilles tendon cross sectional area.

**Figure 4.**
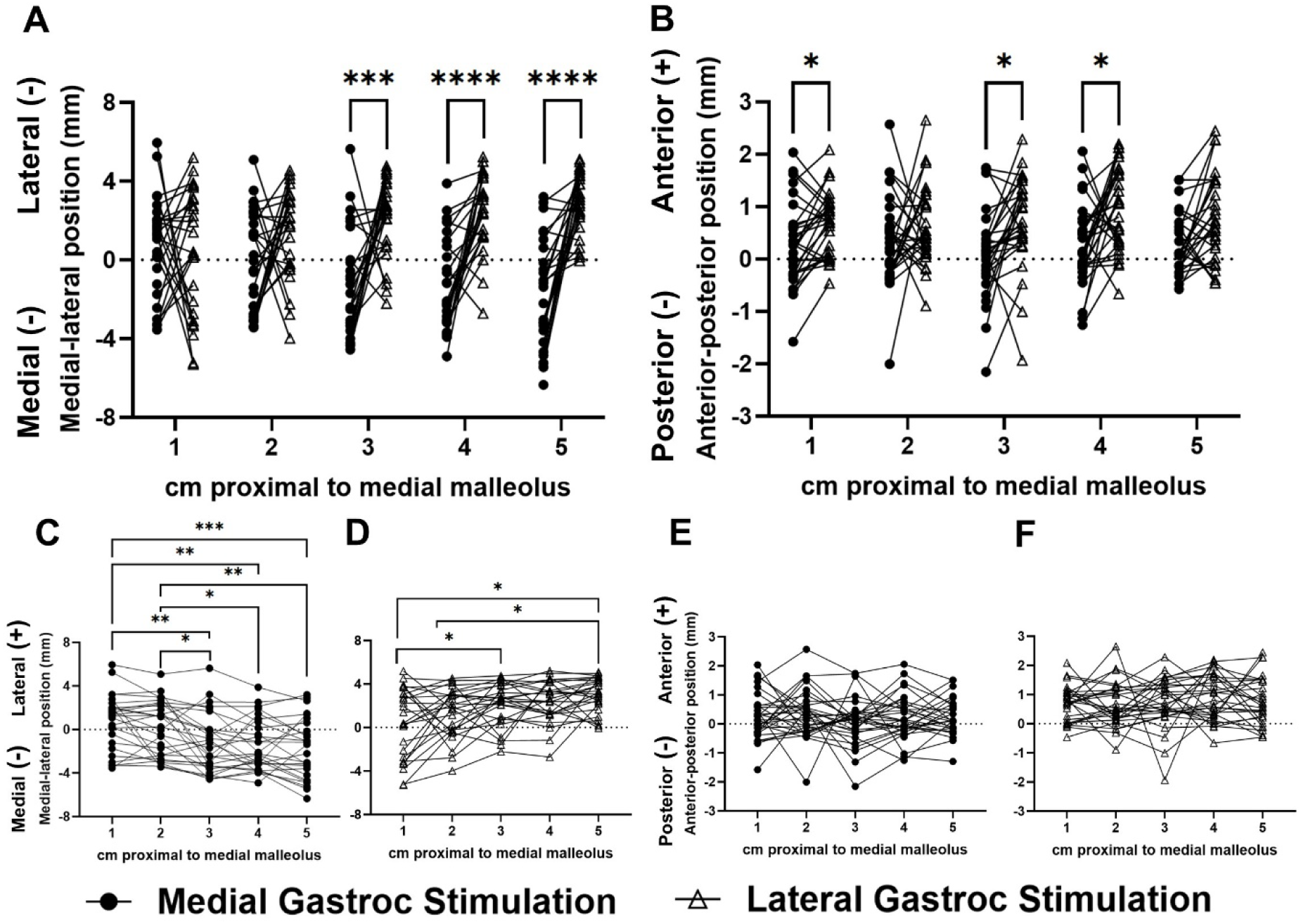
A) Medial-lateral position of the centroid of the most displaced cluster during GM and GL stimulations. B) Anterior-posterior position of the centroid of the most displaced cluster during GM and GL stimulations. C-D) Separate views of the cluster medial-lateral positions during GM and GL stimulations. E-F) Separate views of the cluster anterior-posterior positions during GM and GL stimulations. *p<0.05, **p<0.01, ***p<0.001, ***p<0.0001.

**Figure 5.**
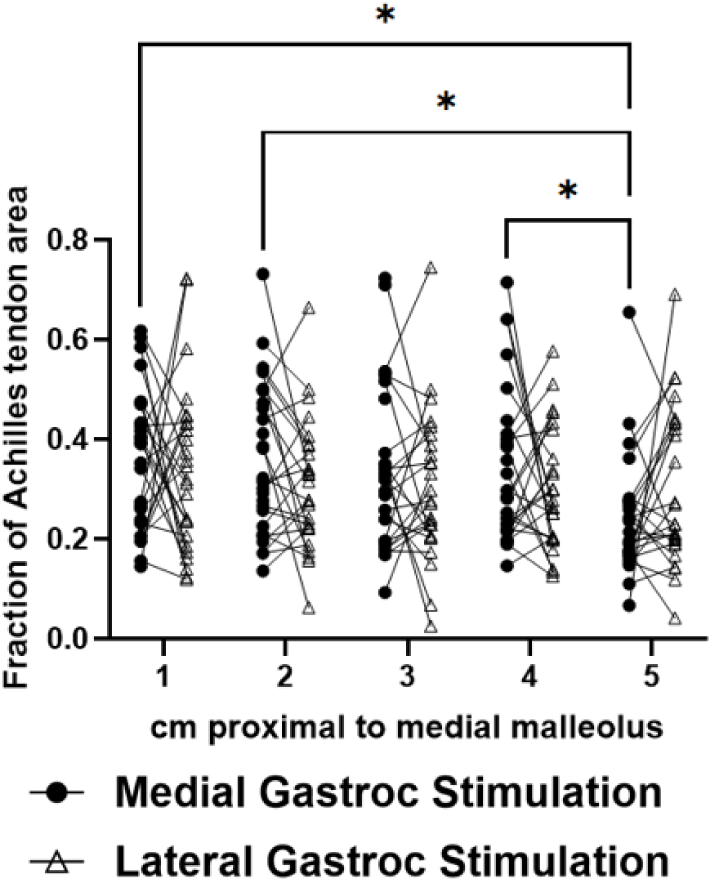
Area of the cluster representing the most displaced region within the tendon cross section, represented as a fraction of the tendon cross sectional area, measured at five locations along the free tendon during both GM and GL stimulations. *p<0.05.

### Bilateral Symmetry

Subtendon cross-sectional area, width, and thickness were not significantly different between right and left sides (**Figure 6**). There were no differences in cluster centroid locations between right and left legs for GL or GM stimulations, and no significant differences in cluster area fractions between right and left sides for any stimulation condition (**Figure 7**).

**Figure 6.**
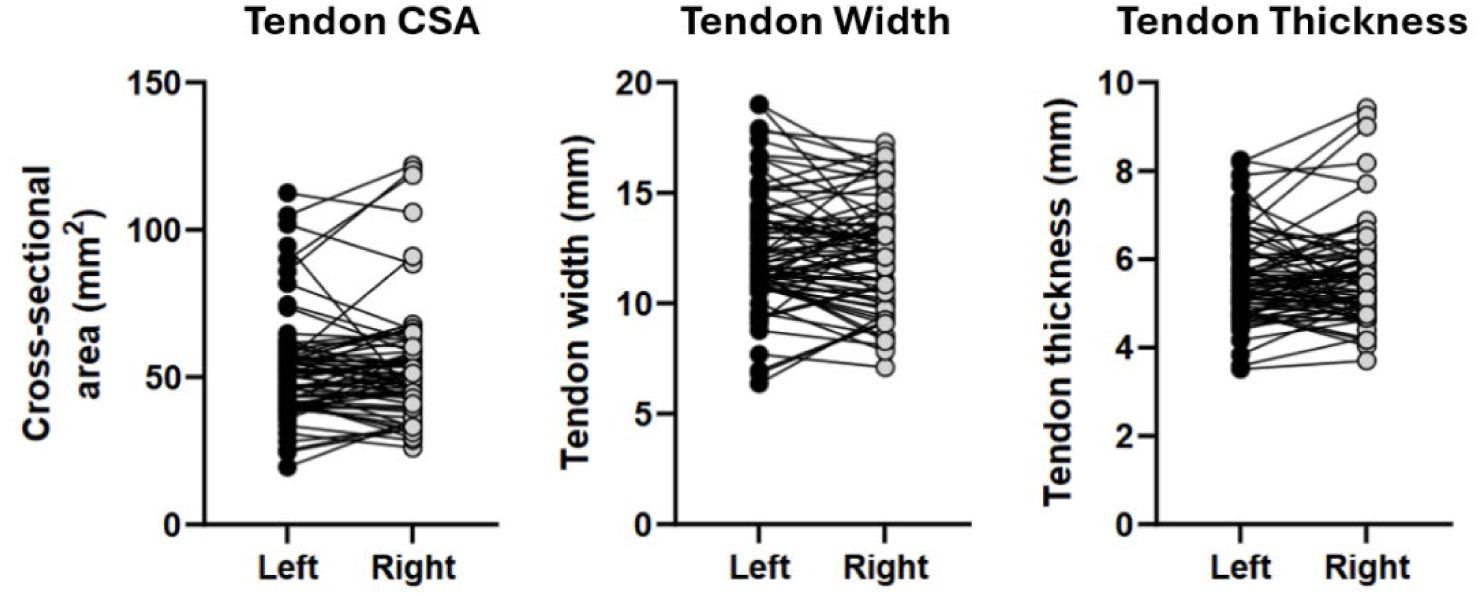
Achilles tendon cross-sectional area, width, and thickness did not differ between right and left legs.

**Figure 7.**
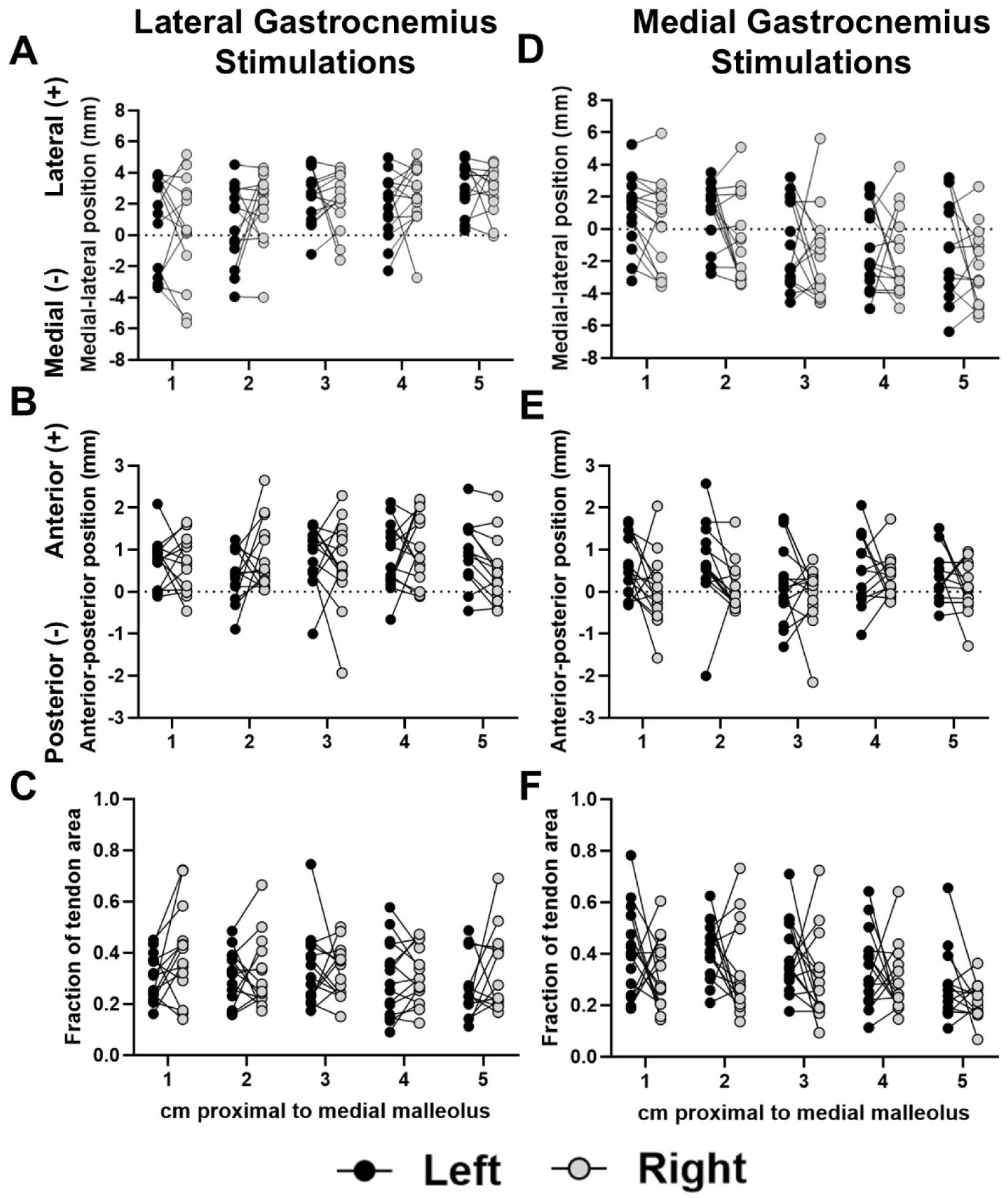
Measurements of cluster centroid positions and area fractions did not differ between right and left legs. Panels A, B, and C display bilateral comparisons of cluster medial-lateral position, anterior-posterior position, and cluster area fraction, respectively, during GL stimulations. Panels D, E, and F display bilateral comparisons of cluster medial-lateral position, anterior-posterior position, and cluster area fraction, respectively, during GM stimulations.

### Reliability

This tool demonstrated high inter-session reliability (ICC>0.83) in identifying cluster locations in response to GL and GM stimulations with the ankle in a neutral position, and moderate inter-session reliability (ICC>0.47) in 20°plantar flexion (**Table 2**).

**Table 2.** ICC values for cluster locations (relative to the Achilles tendon centroid) during GL and GM stimulations in neutral and plantar flexion.

|  | **Lateral Gastrocnemius Stimulations** |  | **Medial Gastrocnemius Stimulations** |  |
| --- | --- | --- | --- | --- |
| **Outcome Measure** | **Neutral** | **Plantar Flexion** | **Neutral** | **Plantar Flexion** |
| **Cluster Medial/Lateral Position** | 0.878 | 0.736 | 0.831 | 0.472 |
| **Cluster Anterior/Posterior Position** | 0.945 | 0.870 | 0.854 | 0.749 |

## Discussion

The purpose of this study was to develop an *in vivo* method to characterize both Achilles subtendon function and morphology within the tendon cross section and along the free tendon. We found significant differences in subtendon displacement patterns between stimulated muscles and between proximal and distal portions of the free tendon. Additionally, we found bilaterally symmetric subtendon behavior in response to individual muscle stimulations in young, healthy adults. Finally, our tool demonstrated excellent repeatability with the ankle fixed in a neutral position. Together, these results support our initial hypotheses.

Our data support our first hypothesis that the Achilles tendon responds differently to individual muscle stimulations along its length and cross section. This hypothesis is supported by the differences in cluster position along the free tendon during GM and GL stimulations, as well as the differences in cluster position within the tendon cross section depending on the stimulated muscle. Moreover, the tissue experienced differential displacement patterns depending on the stimulated muscle, with displacement directions differing significantly within the tendon cross section. Overall, these differential displacement patterns reflect the semi-independent behavior of healthy individual subtendons and suggest that the output from this tool may reflect subtendon function.

In particular, the kinematic data from this study present a new perspective on *in vivo* subtendon function and agree with prior findings. The observed displacements primarily differed in the medial-lateral direction compared to anterior-posterior (**Figure 2**), likely due to the anatomical positions of the GM and GL muscles. Stenroth et al. also found greater nonuniformity of the Achilles tendon during loading in the coronal plane compared to the sagittal (19), corroborating the tendon’s medial-lateral heterogeneity seen in our results. Prior *in silico* models have also observed differences in medial-lateral stress distributions, suggesting that this heterogeneous loading may be a product of Achilles tendon twisting (36). To identify more anterior-posterior heterogeneity, future work should incorporate soleus stimulations as well as stimulations of muscle pairs to investigate the effects of different muscle activations on localized tendon movement. Such work would provide more insight into the load sharing properties between individual subtendons. We also observed greater cumulative displacement values in the proximal region of the free tendon compared to the distal portion, particularly during GL stimulations (**Figure 3**). Prior work reported differential strains within the free tendon compared to the calcaneal insertion (37) that suggest fewer interfascicular connections between subtendons in the proximal free tendon compared to near the insertion site. Overall, these functional differences within the Achilles tendon highlight the need for tools like ours quantifying tendon function in individuals with altered tissue mechanics such as those in aging or injured populations.

Load sharing between adjacent subtendons likely influenced our results. Although lateral load transfer between adjacent tendon fascicles is small at low loads (38), the interfascicular matrix still creates a small amount of friction between subtendons, allowing stimulated muscle’s subtendon to “pull” on its neighboring tissue. Similar effects have been observed in rats; for example, Finni et al. found that activation of a single rat triceps surae muscle caused similar displacements but significantly different strains in adjacent subtendons (39). Moreover, load may transfer differently between the GL and GM subtendons, possibly due to differences in interfascicular matrix properties or myofascial force transmission (40–44). As such, the point clusters identified in the present study likely encompass both the subtendon of interest as well as the adjacent tissue bound by interfasicular matrix. Differing tendon responses to GL and GM stimulations may also result from inherent differences in muscle physiology. For example, the GL has longer fascicles, while the GM has larger pennation angles (11), meaning that similar magnitudes of muscle contractions impart different loads that plantarflex the ankle. While this tool does not capture defined subtendon boundaries, it captures load sharing between adjacent subtendons that we expect will serve as a future biomarker of tendon health. As interfascicular sliding is reduced during injury due to interfascicular adhesions or fibrosis, relative movement between clusters may indicate tendon healing status post-injury and is a subject of future research.

Although this tool is focused on quantifying load sharing between subtendons, our results are similar to prior work examining subtendon morphology. For example, Klaiber et al. found that GL and GM subtendon areas were 15% and 27% of the Achilles tendon cross section, respectively (13). A recent study by Finni et al. employed similar methods and found GL and GM subtendon areas to be 24% and 35% of the tendon cross section, respectively, when measured using similar methods of combined NMES and transverse plane ultrasound imaging (14). Cone et al. found GL and GM subtendon areas to be 41% and 37%, respectively, based on segmentation from high-field MRI (12). Pekala et al. quantified subtendon area at the calcaneal insertion in cadaveric specimens and found the GL subtendon comprising 44% and the GM subtendon comprising 28% of the tendon (4). In the present study, we found the cluster of interest to be 33±16% and 32±14% of the Achilles tendon cross-sectional area during GM and GL stimulations, respectively, which fall within the range of previously reported values. The present study and past work only evaluated healthy, semi-independent subtendons, so localized movement within the tendon cross section is more likely to reflect subtendon morphology. However, in an injured AT, where load sharing is increased because of tendon adhesions and fibrotic scar, we expect this method to be more reflective of function rather than structure. Finally, we also note that the GL and GM relative areas do not sum to 100%, indicating that additional stimulation of the soleus in healthy tendons will further reveal structural information within the tendon cross section.

Although the cluster areas we identified are similar to prior findings, their positions within the Achilles tendon cross section differ from previous work. A prior study by Klaiber et al. employed similar methods of isolated GL and GM stimulations to the present study and found that the GL subtendon was located 4.4 mm lateral and 1.2 mm posterior to the Achilles tendon geometrical center point, while the GM subtendon was located 3.6 mm medial and 1.8 mm anterior (13). In our study, we found that the cluster of interest was located 1.32±2.71 mm medial and 0.09±0.83 mm anterior to the tendon centroid during GM stimulations, and 2.14±1.91 mm lateral and 0.73±0.88 mm anterior to the centroid during GL stimulations when imaged in the midportion of the free AT. Based on prior *in vivo* and cadaveric findings of the deep (anterior) portion of the tendon corresponding to the soleus subtendon (1,2,4,14), we would expect the GL and GM subtendons to be located more posteriorly, particularly in the proximal regions of the free tendon. These discrepancies may be due to differences in data processing methods. For example, Klaiber et al. manually defined each subtendon region of interest based on when the detected displacement vectors first built one convex connected region. Finni et al. also applied similar imaging and NMES methods to ours but found the GM subtendon to be located more posteriorly when manually defining subtendons based on regional echogenicity changes (14). In contrast, our protocol applied unsupervised k-means and density-based clustering to define the clusters of interest. As a result, our final clusters do not necessarily represent a convex region which may alter the resulting centroid position. This approach is advantageous because it is robust to researcher bias toward relative subtendon locations. Moreover, we demonstrated high repeatability in this approach. To our knowledge, only one prior study by Finni et al. tested a similar method for reliability. Moreover, compared to Finni et al., our study evaluates a higher sample size for test-retest reliability and does so using bilateral measurements. These results make this method viable for longitudinal monitoring of patients throughout injury recovery.

Our second hypothesis that the Achilles tendon would display bilateral symmetry was also supported by our results. We found no differences in cluster locations within the tendon cross section between right and left sides. We chose to compare cluster locations relative to the tendon centroid to reflect tendon morphology rather than point kinematics because the stimulation parameters for each muscle were not necessarily equivalent on both right and left legs. These differences may be due to the subject’s activity prior to the testing session as well as slight differences in electrode placement between test days. Given that the k-means clustering algorithm was independent of point location within the tendon cross section, these findings are reflective of inherent anatomical symmetry. These results agree with previous cadaveric studies which did not find bilateral differences in subtendon morphology or Achilles tendon twist types (4). The symmetrical behavior of Achilles subtendons is also an important clinical finding, as it allows the contralateral healthy limb to be used as a reference for altered subtendon behavior after injury.

The findings from this study also confirmed our third hypothesis that this method reliably identifies *in vivo* subtendon behavior across separate testing sessions. The moderate to high ICC values in both neutral and plantar flexion demonstrate that this tool repeatedly captures the expected tendon behavior in response to external stimulation at different levels of tension. As such, future work may utilize this tool to identify altered tendon function due to lengthening post-rupture or increased strain due to gastrocnemius contracture (45,46). We note the higher repeatability (ICC>0.83) for cluster location when the ankle was fixed in a neutral position compared to 20° plantar flexion. This result is likely due to the decreased tension within the tendon while in plantar flexion, resulting in greater displacements in response to the same stimulation amplitude. We estimated that Achilles tendon loading (assuming a 5 cm moment arm (29)) is 0. 0.939±1.726 Nm using our dynamometer measurements of the ankle held neutral position. This load is 95% less than the surgical repair strength of an Achilles tendon rupture repair surgery (30) and should be considered a safe load for patients who are cleared for protected weight bearing in an immobilizing boot. Higher displacements result in more out-of-frame motion and increased tracking errors, so we expect to achieve better repeatability by decreasing the stimulation amplitude when imaging in plantar flexion. We also made the decision to compare cluster positions rather than areas when evaluating repeatability, as cluster area is more sensitive to differential load sharing between subtendons and thus affected by differing stimulation amplitudes between sessions. However, the repeatability of this method makes it viable for quantifying subtendon function *in vivo* in longitudinal studies of tendon healing.

This study has several limitations. First, axial imaging of the Achilles tendon is subject to out-of-plane motion which may affect point tracking results. We chose to use point tracking over speckle tracking, another method commonly used to track motion in ultrasound imaging, as speckle tracking is highly sensitive to out-of-plane motion (33). However, point tracking does begin to fail at large and rapid tendon displacements. We controlled for large out-of-plane displacements by reducing the magnitude of muscle stimulations so as not to induce plantar flexion. However, some subjects experienced greater relative tendon motion in response to muscle stimulation due to individual sensitivities, resulting in failure of the tracking algorithm to converge. As a result, we removed 10 data points out of 300 (15 subjects x 2 legs x 2 muscles x 5 imaging locations) from the final analysis due to such tracking errors.

Using a more gradual stimulation paradigm that achieves the same muscle contraction amplitude over a longer period will increase the number of ultrasound frames with unique tendon displacements. However, our reliability metrics are very good so changing the stimulation paradigm is likely to have diminishing returns. Additionally, we only stimulated the GL and GM muscles of healthy young adults, limiting our understanding of how stimulation of the soleus muscle or altered tissue structure in injured or aging tendons may alter displacement patterns within the tendon cross section. Future work will incorporate isolated stimulation of all three triceps surae muscles and will evaluate individuals with Achilles injuries. Doing so will reveal whether stimulating more muscles is needed to more effectively elucidate subtendon morphology and quantify altered subtendon load sharing.

In conclusion, this study developed a reliable method to characterize Achilles subtendon behavior in healthy subjects using combined NMES and ultrasound imaging in the transverse plane. Our results add to prior evidence of heterogeneity within the tendon in response to isolated muscle contractions. Future work can apply this method to assess both healthy and injured tendons. Such work may provide biomarkers of Achilles tendon function as well as further explore its applications in clarifying subtendon structure *in vivo*.

## Supplemental Material

Supplemental Figure S1: https://doi.org/10.6084/m9.figshare.29189753.v1

## Data Availability

All data from this study are available from the corresponding author upon request.

## Grants

This study was supported by NIH/NIAMS P50AR080581 (JRB) and NSF Grant DGE-2236662 (KSS)

## Disclaimers

The authors have no disclosures, financial or otherwise.

## Author Contributions

Conceived and designed research (KSS, TJH, JRB); Analyzed data (KSS); Performed experiments (KSS); Interpreted results of experiments (KSS); Prepared figures (KSS); Drafted manuscript (KSS); Edited and revised the manuscript (KSS, JRB, TJH); Approved final version of the manuscript (JRB)

## Acknowledgements

We thank Maggie Wagner for assistance in EMG data collection and processing. Figure illustrations created with BioRender.com.

