## Supplementary material for "In vivo characterization of Achilles subtendon function and morphology within the tendon cross section and along the free tendon": Figure S1

**Figure S1.** Fascicle tracking trials used to confirm isolated recruitment of a single gastrocnemius muscle. Fascicle length tracked using a previously validated automated tracking algorithm. Figure shows representative tracking frames of the GM muscle at rest (A) and during the stimulation (B). Plots show quantified fascicle lengths of the stimulated and non-stimulated muscles during lateral gastrocnemius (GL) (C) and medial gastrocnemius (GM) (D) stimulations. Fascicles shortened more on the stimulated muscle compared to the resting length compared to the non-stimulated muscle. Data represents results from a single trial.

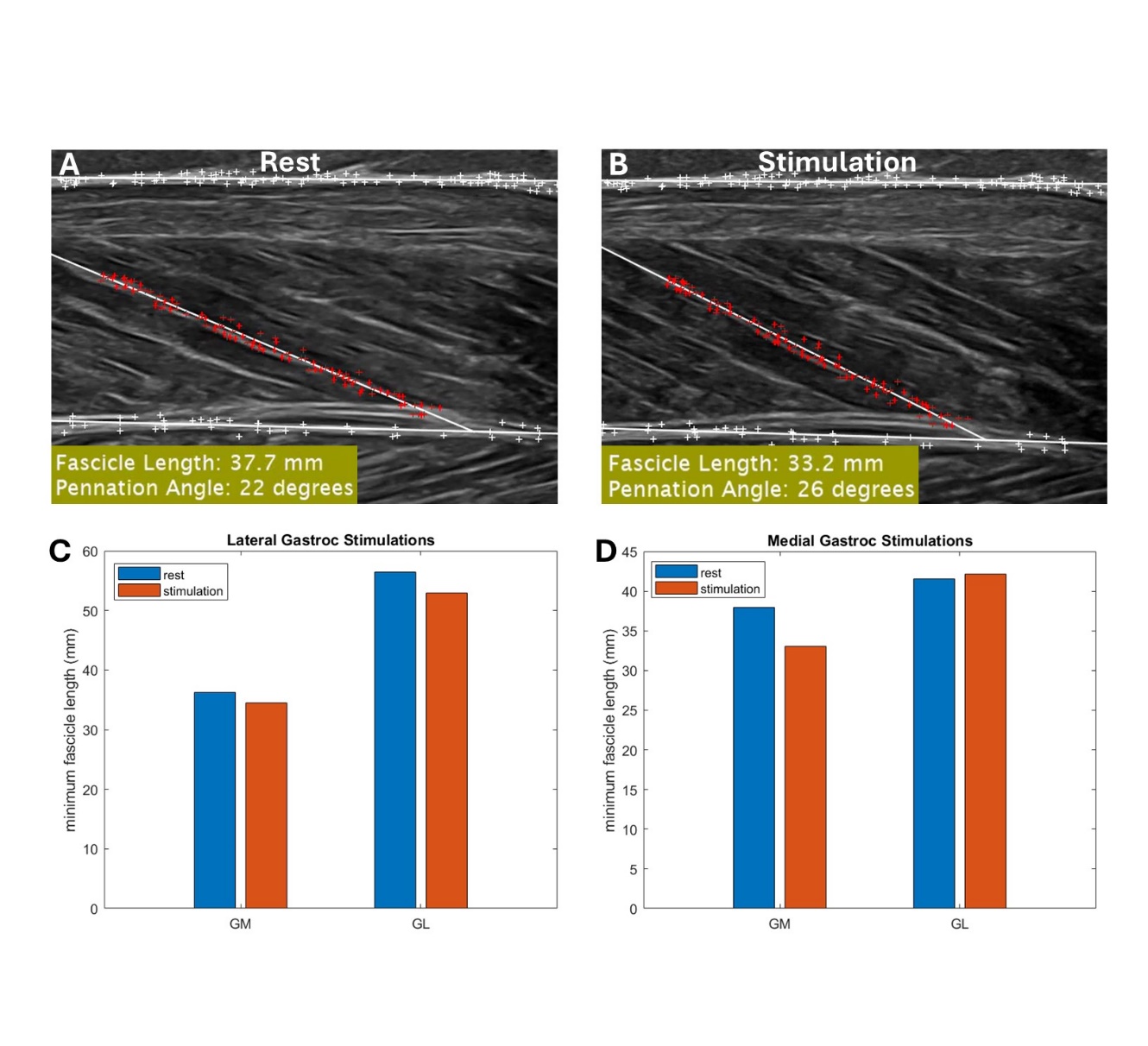
